# Pleiotropy links host axes of defense against a pathogen

**DOI:** 10.64898/2026.10.12.758686

**Authors:** Michael Blazanin, Nowmita Nasir, Emilie Haniak, Bardia Masudy, Phae Elton, Sarah P. Otto, Kayla C. King

## Abstract

Organisms face ubiquitous threats from pathogens, and in response hosts have evolved a cascade of defenses to avoid, resist, or tolerate infection. While these stepwise defenses may act independently, they can be pleiotropically linked via shared molecular mechanisms, regulation, and crosstalk. Here, we tested the extent to which pleiotropy shapes multiple axes of defense using knockouts of 16 single genes involved in innate immunity of *Caenorhabditis elegans* against the opportunistic bacterial pathogen *Pseudomonas aeruginosa*. Host genotype had variable effects on components of host defense. Traits such as infection-induced mortality and body size varied frequently across host genotypes, while behavioral avoidance, bacterial uptake, and susceptibility to bacterial colonization varied less frequently. Broadly, pleiotropy between defenses was common, driving patterns of variation that linked several traits together. However, important components of defense like mortality, avoidance, and susceptibility to colonization were more independent from one another. Our results show that pleiotropy between defenses can be common, highlighting the importance of moving beyond single phenotypes to understand multiple dimensions of host-pathogen interactions.

## Introduction

Across the tree of life, organisms face ubiquitous threats from pathogens (Schmid-Hempel 2021). In response, hosts have evolved numerous defenses to avoid, resist, or tolerate infection. Defenses often form a cascade-like process over the course of an individual infection (Hall et al. 2017). Early defenses include avoidance, which pre-emptively reduce the exposure of hosts to pathogens (Gibson and Amoroso 2022), as well as defenses against the initial establishment of the infection. Once the pathogen is established, hosts often resist the growth of the pathogen. Additionally, hosts may have mechanisms to tolerate infection (Medzhitov et al. 2012; Pagán and García-Arenal 2020) or that alter onwards transmission of the pathogen (Duffy 2021).

While these stepwise defenses might act independently, they might also be linked via shared genetic mechanisms that pleiotropically alter multiple components of defense. Although we often have a good understanding of individual components of host defense, our understanding of the importance of pleiotropy across different host defenses is much more limited (Hall et al. 2017; Mommer et al. 2026). Pleiotropy that links different components of host defense could have important implications for disease dynamics and evolution. For instance, pleiotropy shapes the genetic variation in traits available for natural selection. Available variation biases the direction of evolution (Schluter 1996; Mallard et al. 2023), constraining change against trade-offs, and accelerating change along trade-ups. These relationships can potentially allow selection to efficiently shape multiple traits simultaneously (Wessinger and Hileman 2016).

Here, we address the gap in our knowledge of pleiotropy across host defenses by exposing the model eukaryote *Caenorhabditis elegans* to an opportunistic bacterial pathogen. This nematode exhibits a diversity of defenses against bacterial pathogens, including several conserved pathways shared with other eukaryotes (Kumar et al. 2020). *C. elegans* has both innate and learned avoidance behaviors towards bacterial pathogens that reduce its exposure to harmful bacteria while foraging in decaying organic matter (Meisel and Kim 2014). The nematode also has physical barriers that limit exposure to pathogens, including a grinder in the pharynx that prevents intact bacteria from reaching the gut (Avery and You 2012). Finally, several signaling pathways play central roles during the nematode’s innate immune response to infection. These pathways include the insulin/IGF-1-like signaling pathway (Murphy and Hu 2018), the p38 MAP Kinase pathway (Kim et al. 2002), the Dar phenotype (Hodgkin et al. 2000), and several signaling pathways in the TGF-β family (Yamamoto and Savage-Dunn 2023), including the DBL-1/BMP-like, the DAF-7, and the TIG-2/BMP-like TGF-β/Activin-like ligand TIG-3 pathways. These innate immune pathways can modulate the suppression and tolerance to an infection.

We tested for the importance of pleiotropy across distinct components of host defense using knockouts of 16 innate immunity genes. These genes have been previously implicated in one or more of *C. elegans* defenses against infection by the bacterial pathogen *Pseudomonas aeruginosa*, covering all the major known pathways of defense. *P. aeruginosa* is a widespread opportunistic pathogen which uses conserved mechanisms to infect animals, including humans and *C. elegans* (Hajdú et al. 2024). The pathogen enters *C. elegans* through the normal bacteria-eating behavior of the nematode, then takes up residence in the gut, establishing an infection that ultimately leads to death of the host after 3-4 days (Tan et al. 1999). We show that immunity genes frequently have pleiotropic effects. We quantified several components of host defense, including avoidance, uptake, susceptibility to colonization, and mortality. We found that genotype had variable effects on host defenses, altering some traits more frequently and strongly than others. Nevertheless, pleiotropy across distinct components of defense was common, with many genes altering multiple defenses. Additionally, we discovered fairly strong patterns of covariation that link some host defenses together while other defenses remain independent.

## Methods

### Strains and growth conditions

In order to carry out multiple assays, we maintained working stocks of actively growing *Caenorhabditis elegans* populations at 20 °C on 90mm Nematode Growth Medium plates [NGM: 3 g/L NaCl, 17 g/L agar, 2.5 g/L peptone (Millipore #70169), 5 mg/L cholesterol, 1mM CaCl2, 1 mM MgSO4, 21.7 mM KH2PO4, 3.3 mM K2HPO4] with a lawn of *Escherichia coli* OP50 bacteria as food. All *C. elegans* strains were acquired from the Caenorhabditis Genetics Center (CGC) at the University of Minnesota (Table 1). Long-term worm stocks were stored at -70 °C in worm freezing solution (300 g/L glycerol, 5.86 g/L NaCl, 6.8 g/L KH2PO4, 5.6 mM NaOH, 3 mM MgSO4). Working lines of *C. elegans* were passaged once per week onto fresh lawns of *E. coli* and re-thawed from frozen at most every 10 passages to limit host evolution. Bacterial stocks were stored at -70 °C in 25% glycerol. Working bacterial stocks were struck from frozen onto LB (10 g/L tryptone, 5 g/L yeast extract, 10 g/L NaCl) plates (15 g/L agar) and grown until colonies formed, then stored at 4 °C. Lawns were created by inoculating a bacterial culture from a single colony into ∼25 mL LB liquid in a 50 mL Falcon tube, growing overnight shaking at 30 °C, then spreading 100 µL and incubating at 30 °C overnight.

**Table 1.** Strains used in this study.

| Strain | Genotype |
| --- | --- |
| *P. aeruginosa* PA14 (pathogen) | Wild-type |
| *E. coli* OP50 (food) | Wild-type |
| *C. elegans* N2 | Wild-type |
| *C. elegans* TJ1052 | age-1(hx546) II. |
| *C. elegans* RB2430 | tig-2(ok3336) V. |
| *C. elegans* DA1814 | ser-1(ok345) X. |
| *C. elegans* CB7007 | llys-3(ok3222) IV. |
| *C. elegans* VC3044 | dbl-1(ok3749) V. |
| *C. elegans* RB2302 | daf-7(ok3125) III. |
| *C. elegans* KU25 | pmk-1(km25) IV. |
| *C. elegans* FX1053 | ins-11(tm1053) II. |
| *C. elegans* CF1038 | daf-16(mu86) I. |
| *C. elegans* DA465 | eat-2(ad465) II. |
| *C. elegans* RB911 | fshr-1(ok778) V. |
| *C. elegans* VC3056 | zip-2(ok3730) III. |
| *C. elegans* SJ17 | xbp-1(zc12) III; zcls4 V. |
| *C. elegans* RB1330 | npr-1(ok1447) X. |
| *C. elegans* CX13079 | octr-1(ok371) X. |
| *C. elegans* RB1321 | C56G3.1(ok1439) X. [npr-8 loss-of-function] |

### Exposing hosts to bacteria

Our methods for exposure of *C. elegans* to bacteria (below) were modified from those in the literature, which vary considerably from paper to paper (Table S1). *P. aeruginosa* was sourced from Kevin Foster (Oxford University). Nematodes were age-synchronized by bleach-washing (2% hypochlorite solution, 1.67M NaOH for 6-10 minutes) gravid hermaphrodites then hatching larval stage 1 (L1) larvae in M9 buffer (3 g/L KH2PO4, 6 g/L Na2HPO4, 5 g/L NaCl, 1 mM MgSO4) overnight. L1 nematodes were then grown on food plates for 2 days before being transferred to exposure conditions. To create exposure plates, food (*E. coli*) or pathogen (*P. aeruginosa*) bacteria were inoculated from a single colony into 5 mL LB liquid in a 50 mL Falcon tube, then grown overnight shaking at 30 °C. Bacterial cultures were then normalized to an OD600 of 0.05 (food: ∼1.7×10^8^ CFU/mL, pathogen: ∼6.2×10^7^ CFU/mL), and 100 µL was spread onto 60 mm plates of ‘NG’ media (same as NGM, but with 3.5 g/L peptone), using an L-shaped spreader to ensure there were no gaps in the lawn. Plates were incubated at 30 °C for one day then 25 °C for one day. After ∼48 hours we transferred 100 L4/young adult nematodes to each exposure plate and incubated at 25 °C.

### Quantifying host survival from pathogen exposure and bacterial load in the gut

For survival and bacterial load, three replicate plates (for survival) or three for each timepoint (for load) were exposed to food (as a control) or pathogen bacteria, and data were collected at 0, 24, 48, 72, and 96 hours. We did not collect data from control plates at 96 hours because the next generation of nematodes were already reaching adulthood.

To quantify survival, the number of adult living worms present on each plate was counted, poking worms as necessary with a worm pick to observe movement to determine survival. The area under the curve of the number of adult worms relative to the initial number (Fig S4) was calculated for each replicate for the first 72 hours of exposure then divided by 3 (the maximum possible AUC). dbl-1 on control plates was not counted at 72 hours because the next generation of nematodes had reached adulthood, so the number of adult worms relative to initial was treated as 1 for the purposes of AUC calculation.

To quantify the population size of gut bacteria, we used methods modified from Hoang et al. (Hoang et al. 2024). At each timepoint, 10 living adult worms (when possible) were picked off each of three replicate plates into 498 µL of 4 °C chilled M9 buffer with 0.01% Triton-x on ice. Worms were not picked for pathogen densities at 0 hours, as they had not yet been exposed to the pathogen. After picking, worms were chilled at 4 °C on ice for 30 minutes to stop peristalsis. Then 2 µL of a hypochlorite solution was added to reach a final concentration of 0.0012% hypochlorite (0.02% commercial bleach). Worms were then stored on ice for 10 minutes to kill external bacteria. Worms were washed twice with chilled M9 buffer and homogenized by bead beating at 30/s for 5 minutes. Bacteria were serially diluted in M9 and spotted onto LB plates, which were incubated until colonies could be counted and the colony-forming units per worm calculated.

### Quantifying host avoidance of bacteria

To quantify worm avoidance of bacteria, we used a lawn-leaving assay modified from (Amoroso et al. 2024, 2025). Three replicate exposure plates for each nematode strain and bacterium were prepared as described above except that bacterial cultures were normalized to an OD600 of 0.3 (food: ∼1.0×10^9^ CFU/mL, pathogen: ∼3.7×10^8^ CFU/mL), and 50 mL was spread onto 60 mm plates of NG media into a circular area in the center of the plate with a 24.5 mm diameter. After 48 hours incubation (at 30 °C and 25 °C as described above), 20 worms were added to the center of each plate. The number of living adult worms on and off the bacterial spot were counted at 0, 24, and 48 hours after plating.

### Quantifying host morphology

To quantify differences in nematode size and shape across development, nematodes were age synchronized and plated onto one food plate per strain as described above for exposure. After 24, 48, 72, 96, 120, 144, and 188 hours, at least three worms were recorded at 10 frames per second for up to one minute using a Leica M205 FA microscope with a K5C camera. As needed, adult worms were picked onto a fresh food plate to ensure the original adults did not become indistinguishable from developing larva. Videos were analyzed per worm with WormLab v2024 (MBF Bioscience) with length fitting, detection frequency every 1 frame, back-tracking, and manually set values for threshold, background smoothing, fill holes, position tolerance, and shape tolerance. Worm width, length, projected body area, and fit were exported from Wormlab, but fit did not always reflect the best-fitting frames, so in some cases best-fitting frames were manually marked. Then the data from the best-fitting frame(s) for each worm were averaged together.

### Quantifying host uptake of bacteria

To approximate rates of host uptake of bacteria, we quantified rates of host pharyngeal pumping on food and pathogen bacterial exposure plates. We exposed three replicates of each host strain to each of food or pathogen bacteria. Videos at least 30 seconds long were taken to record pharyngeal pumping motion, with T0 videos taken right after exposure, and T1 videos taken after 24 hours exposure at 25 ⁰C. Videos were recorded using Leica M205 FA microscope with a K5C camera. Muscle movements denoting each pharyngeal pump were counted manually from each video.

### Analysis and batching design

For experimental tractability, each trait was measured separately, with strains split into 3 to 5 batches per trait. Wild type (N2) worms were included in every batch. Batch-correction was carried out by fitting a linear mixed-effects model with lme4 (Bates et al. 2015) of only the wild type data, calculating random batch effects, then subtracting these effects from all data. For bacterial density and pharyngeal pumping this was carried out for each timepoint. Some data deviated from this slightly: for mortality and avoidance we used a generalized model with a binomial family fitted at each timepoint, and for mortality AUC we used a generalized model with a beta family after calculating AUC from raw data. There was only one batch of WormLab data.

Statistical comparisons to wild type were carried out by fitting a linear model of batch-corrected values (AUC of survival, avoidance, log10-transformed bacterial density, pumping rates, or area) as a function of genotype and bacteria, then using estimated marginal means contrasts of each genotype against N2 with emmeans (Lenth 2025). For pumping data, some timepoints had more replicates than others; to avoid any bias in which worms were recorded, and because replicates were not linked across timepoints, we randomly sampled replicates from each timepoint to form complete sets, then carried out statistics.

Clustering of traits and genotypes (Fig 2) was done by mean-centering and scaling the traits, then clustering each of traits and genotypes with Euclidean distance and Ward’s method for agglomeration (CITE). Analysis of correlations between traits (Fig 3) was carried out by calculating Pearson’s correlation coefficient and its significance between every pair of traits. Clustering of correlations was done by Euclidean distance and complete linkage of the absolute value of correlation coefficient.

## Results

To test for the importance of pleiotropy in host axes of defense against pathogens, we first quantified the effects of host genotype on five traits related to pathogen interaction: behavioral avoidance of bacteria, uptake rates of bacteria, gut bacterial population size, survival when exposed to bacteria, and body size. We found substantial effects of genotype on all five traits (Fig 1), generally finding stronger effects of genotype when hosts interacted with the pathogen than with control food bacteria.

**Figure 1.**
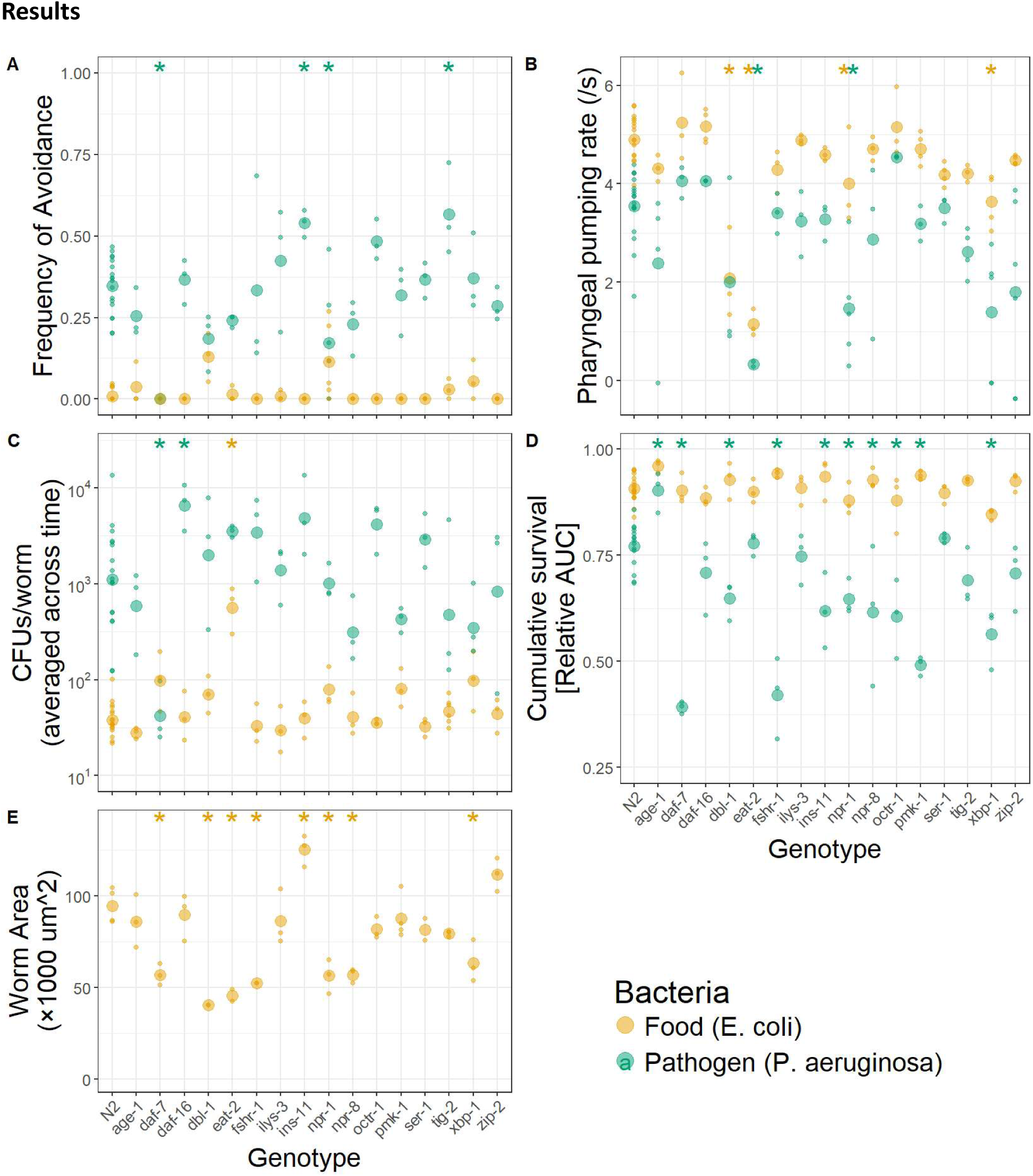
Host genotypes vary in responses to bacteria. Wild-type (N2) and mutant genotypes with single gene knockouts were assayed for five traits: A) behavioral avoidance of bacteria, B) uptake rates of bacteria, C) gut bacterial population size, D) survival when exposed to bacteria, and E) body size. Small points denote individual replicates while large points denote the average, asterisks (at top) denote genotypes that significantly differ (p < 0.05) from wild type. **A.** Hosts were exposed to a spot of food or pathogen bacteria on an otherwise-empty agar plate for 24 hours, then the number of hosts on the spot versus avoiding the spot was counted. **B.** Hosts were exposed to lawns of food or pathogen bacteria for 24 hours, at which point the pharyngeal muscle movements of at least 3 worms per strain were manually counted from 30s videos. **C.** Hosts were exposed to lawns of food or pathogen bacteria for three days. 10 worms of each genotype were picked from each of three replicate plates for each strain in each condition and at each timepoint. After picking, the exterior of the worms was sterilized before the gut contents were plated to determine the average colony-forming units (CFUs) per worm. **D.** Hosts were exposed to lawns of food or pathogen bacteria for three or four days, respectively. Three replicate plates were counted for each genotype in each condition. Plotted is the area under the survival curve, relative to the maximum possible. **E.** Hosts were reared on food bacteria for two days then imaged (n ≥ 3 each genotype). After imaging, the projected body area of each worm was calculated using WormLab.

The first defense hosts have against pathogens is behavioral avoidance. We found that host genotype had minimal effects on avoidance of control food bacteria, which was generally negligible (Fig 1A). In contrast, host genotype had strong and variable effects on avoidance of pathogen bacteria (Fig 1A), including genotypes with reduced ability to avoid the pathogen (daf-7, npr-1) and increased ability to avoid the pathogen (ins-11, tig-2).

Once hosts encounter a pathogen, their next defenses restrict the uptake of the pathogen. In *C. elegans*, many bacteria enter the gut through the normal bacteria-eating behavior of the nematode. As a proxy for uptake rates of bacteria into the gut via feeding, we quantified the rate of pharyngeal pumping contractions. Most host genotypes had minimal effects on feeding rate, which was generally higher on control food bacteria than pathogen bacteria (Fig 1B) and increased through time (Fig S2). However, a couple host genotypes (eat-2 and npr-1) exhibited reduced feeding rates on both food and pathogen bacteria.

Once a host has become infected, its next lines of defense restrict the accumulation and proliferation of a pathogen. We quantified differences in gut bacterial populations, finding that host genotype had minimal effects on gut bacterial density (Fig 1C). Hosts generally had negligible populations of control food bacteria in their gut, with only one genotype (eat-2) significantly increasing densities. Hosts nearly always experienced much more substantial colonization by the pathogen bacteria than control bacteria, but genotype again had minimal effect, with only two genotypes (daf-7, daf-16) significantly differing from wild type.

Once infection is established, host defenses can also control the susceptibility to mortality. We quantified differences in survival, finding that host genotype had substantial effects on mortality (Fig 1D). Hosts experienced minimal mortality on control food bacteria, with no significant effects of genotype. In contrast, hosts experienced substantial levels of mortality when exposed to the pathogen, with half (8/16) of the genotypes significantly differing from wild type, including one genotype that had significantly increased survival, and seven genotypes with significantly decreased survival.

The final trait we quantified is host body size, which could be a potentially important correlate of pathogen-related traits. We found that genotype had significant effects on host size at the young adult stage (when hosts were transferred from rearing to exposure conditions in our experiments), with seven genotypes being significantly smaller than wild type, and one genotype (ins-11) being significantly larger than wild type (Fig 1E).

We explicitly tested for relationships among the traits we had measured. First, we tested how frequently host genotype pleiotropically altered multiple traits, finding that pleiotropy was common (Fig 2). Among the 13 knockout genotypes that exhibited any significant differences from wild type, eight had significant differences from wild type in multiple phenotypes. We also found similar results when traits were analyzed across multiple timepoints (Fig S8). Interestingly, genotypes tended to be fairly unique in their changes, with minimal clusters among genotypes emerging other than three knockouts that had minimal effects and so clustered closely to wild type. Second, we tested for patterns in the correlations between traits. We found one cluster of highly correlated traits (Fig 3): across all genotypes, larger hosts tended to have higher uptake rates of both control food and pathogen bacteria, tended to avoid food bacteria less, and tended to have fewer intact food bacteria in their gut. The remaining traits (survival against control food bacteria, survival against pathogen bacteria, avoidance of the pathogen, and densities of pathogen bacteria in the gut) tended to be only weakly correlated with other traits. We also found similar results when traits were analyzed across multiple timepoints (Fig S9).

**Figure 2.**
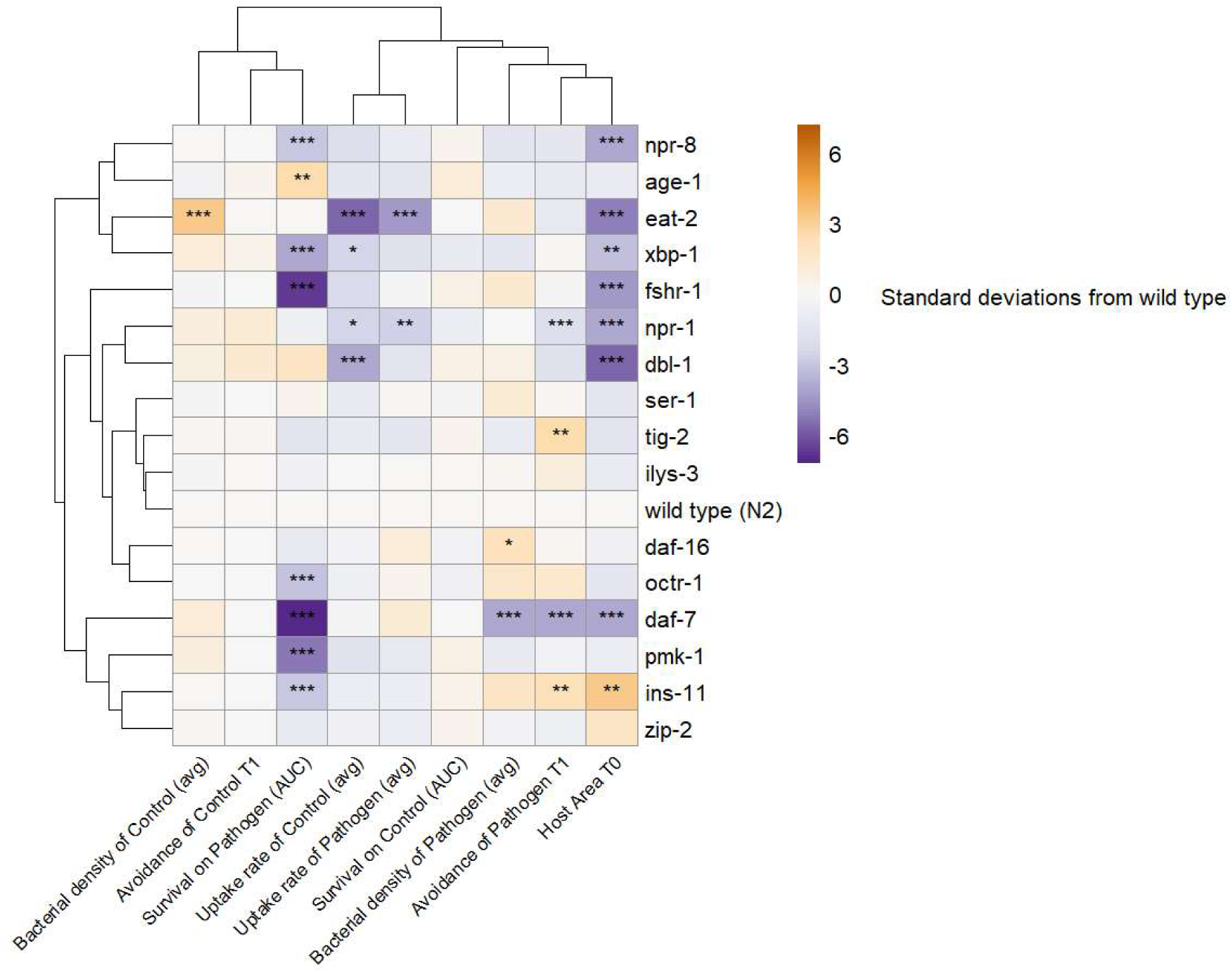
Pleiotropy is common across host traits related to pathogen interactions. Data from Fig 1 was replotted after being centered at the wild type value and scaled proportional to the pooled standard deviation across genotypes. Clustering of traits and genotypes was done by mean-centering and scaling the traits, then clustering either traits or genotypes with Euclidean distance and Ward’s method for agglomeration. Asterisks denote two-tailed significant differences from the wild-type (N2) of estimated marginal means of batch-corrected data: *** for p < 0.001, ** for p < 0.01, and * for p < 0.05.

## Discussion

Hosts have a cascade of defenses against pathogen infection (Hall et al. 2017), yet little is known about genetic linkage across these axes of defense. Here, we used genotypes of *C. elegans* with knockouts of known innate immune genes to characterize how genetic variants altered interactions with the opportunistic bacterial pathogen *P. aeruginosa*. We found that genotype had variable effects on host defenses. Pleiotropy was common, with many single gene knockouts having effects that bridged across multiple components of host defense. Among the traits we measured, we found that several strongly covaried, suggesting that the genetic architecture of the host may coregulate them jointly. In contrast, important axes of defense like mortality, avoidance, and susceptibility to colonization tended not to be consistently correlated with other traits. Ultimately, our results show that components of host defense can be highly independent while still experiencing frequent pleiotropic linkage.

Host-pathogen interactions have often been conceptualized as a series of steps, from exposure to establishment, colonization, and transmission (Hall et al. 2019; Mommer et al. 2026). The distinctions between these stages provide important insights, and inform our ecoimmunological understanding of host-pathogen interactions (Graham et al. 2011). For instance, increased host immunity has been predicted to drive pathogens to evolve increased levels of virulence (harm to their host), but only when immunity operates on colonization or mortality (Gandon et al. 2001). Similarly, modeling the dynamics of pathogen infection across multiple stages can lead to qualitatively different outcomes, because nonlinearities within and between stages of infection can interact (McCallum et al. 2017). For instance, the success of vaccination programs can depend strongly on what stage of immunity is boosted by the vaccine (Gandon et al. 2003). Here, we have attempted to independently quantify pathogen dynamics at each stage of infection in the *C. elegans* model system: measuring avoidance to approximate exposure, uptake as a proxy for establishment rate, colonization dynamics over time, and host mortality for transmission. It is clear that, indeed, pathogen infection dynamics can vary strongly between host genotypes at one stage of infection, but strongly resemble each other at other stage(s) of infection. Thus, our data support the importance of models that distinguish between pathogen dynamics at different stages of infection (Gandon et al. 2001, 2003).

Disparate host defenses can trade-off with one another or other components of host fitness. For instance, a seminal paper found that resistance and tolerance in plant defenses against herbivory tend to trade off (Fineblum and Rausher 1995). Largely, evidence for trade-offs between resistance and tolerance is quite mixed (Mesa et al. 2017; Montes et al. 2020; Pagán and García-Arenal 2020). Our results add to this trend, with survival and pathogen colonization tending to have weak and non-significant relationships with the other traits we quantified. Although, the variation in our dataset reflects loss-of-function mutations and not adapted phenotypes (Fig 3). Other trade-offs can also shape patterns of host defense. In plants, the ‘Leaf Economics Spectrum’ defines a cluster of highly correlated traits that also correlates with resistance against pathogen infection (Welsh et al. 2020), and in animals ‘pace-of-life’ variation tends to drive covarying changes in numerous traits, including susceptibility to disease (Ricklefs and Wikelski 2002). Similarly, we did find a cohort of traits that covaried strongly among our host strains, including size, uptake of bacteria, food-seeking behavior, and levels of intact food bacteria in the gut. Such covariation might be important, since pleiotropy between immune-related traits (e.g., uptake) and non-immune traits (e.g., size) can constrain the evolution of immunity genes in hosts (Williams et al. 2023). Altogether, our data support the understanding that trade-offs can powerfully shape trait covariation.

**Figure 3.**
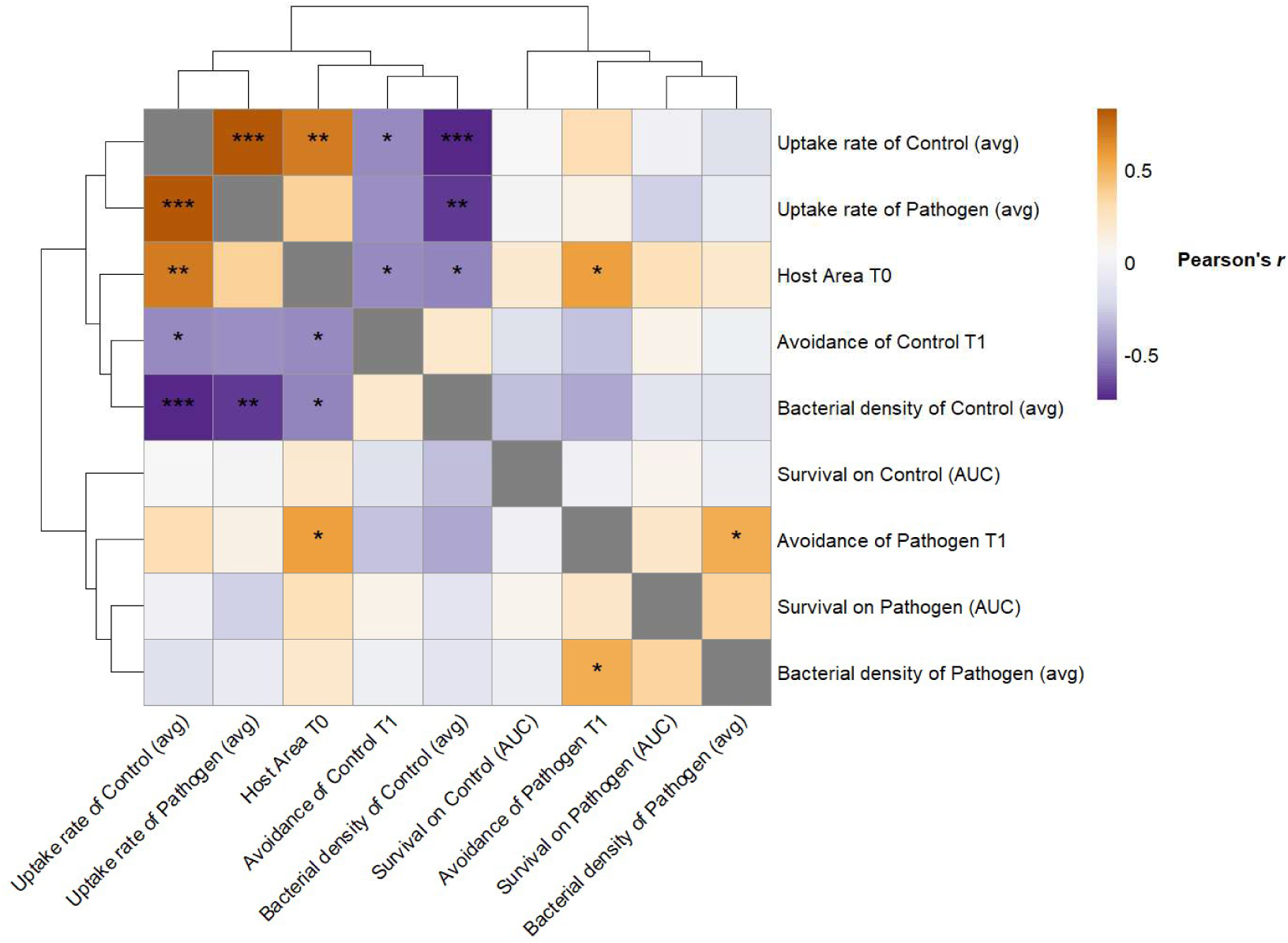
Some traits highly correlate while others are largely independent. Data from Fig 1 was used to calculate the correlation of each trait against each other trait among the 17 host genotypes. Clustering of traits was done by Euclidean distance and complete linkage of the absolute value of the correlation coefficient. Asterisks denote the two-tailed significance of Pearson’s *r*: *** for p < 0.001, ** for p < 0.01, and * for p < 0.05.

While these findings reveal important facets of a model host-pathogen interaction, they also raise new questions. The gene knockouts studied here were chosen because previous work had implicated them as central components to one or more aspects of the immune response to pathogen infection (Kurz and Tan 2004; Irazoqui et al. 2010b; McEwan et al. 2012; Portal-Celhay et al. 2012; Pukkila-Worley and Ausubel 2012; Lee et al. 2018; Williams and Schumacher 2018; Foster et al. 2020; Ciccarelli et al. 2024; Xiao et al. 2024). Further work is needed to test whether the patterns of pleiotropy and covariance we found reflect genome-wide patterns or are specific to central components of the innate immune system. Similarly, more work is needed to collect analogous datasets in other species to draw broader comparative conclusions about the structure of the genetic architecture that governs host-pathogen interactions more broadly (Letendre et al. 2022). At a more specific scale, fine-grained investigations are needed to better understand the dynamics of infection in *C. elegans* as a model system. For instance, bacteria must persist through the pharynx and grinder to establish an infection in the gut, but little is known about how rates of uptake via pumping [∼20 bacteria/pump (Ferris et al. 1997; Schulenburg et al. 2004)] translate to rates of introduction to the gut (Browning et al. 2013; Hsiao et al. 2013). Finally, the discovery of these patterns of host defense raises the question of how such variation in host defenses ultimately shapes their pathogens over longer evolutionary timescales.

Hosts face a myriad of threats from pathogens, and their defenses against these intruders can be layered and complex (Schmid-Hempel 2021). Little is known about the genetic basis of relationships among host defense traits. We have found these traits can exhibit a mixture of strong covariation and independence while pleiotropy commonly links across defenses.

## Acknowledgements

Thanks to Alita Burmeister and Caroline Turner for feedback on the manuscript. MB acknowledges that this material is based upon work supported by the NSF Postdoctoral Research Fellowships in Biology Program under Grant No. 2508854 to M.B. and an NSERC Canada Excellence Research Chair to K.C.K. The funders had no role in this project’s development, design and delivery.

