## Supplemental Materials for "Pleiotropy links host axes of defense against a pathogen"

**Table S1. Methods vary greatly to induce *Pseudomonas* slow-killing of *C. elegans*.** Published papers were reviewed to compare methods, including conditions for: 1) liquid inoculation, 2) plating, 3) pre-exposure plate incubation, and 4) exposure. "O.N." is short for "Overnight". "NGM" media contains 2.5 g/L peptone, while "NG" media (i.e. 'SK' slow-killing media) contains 3.5 g/L peptone.

| Paper | Year | Liq. media | Liq. temp. (time, vol.) | Plated vol. | Plate media | Plate incubation | Expos. temp. |
|---|---|---|---|---|---|---|---|
| (Tan et al. 1999) | 1999 | King's broth | O.N. | 5-10 µL | NG | 37C (24hr), 25C (8-24hr) | 25C |
| (Garsin et al. 2003) | 2003 | ? | ? | ? | ? | ? | ? |
| (Troemel et al. 2006) | 2006 | LB | ? | ? | NG | 37C (24hr), 25C (24hr) | 25C |
| (Powell and Ausubel 2008) | 2008 | LB | 37C (12-16hr, 2-5mL) | 10 µL | NG | 37C (24hr), 25C (24hr) | 25C |
| (Irazoqui et al. 2008) | 2008 | ? | ? | ? | ? +FUDR | ? | ? |
| (Powell et al. 2009) | 2009 | ? | ? | ? | ? +FUDR | ? | 23C |
| (Irazoqui et al. 2010a) | 2010 | LB | ? | ? | NG | 37C (24hr), 25C (24hr) | 25C |
| (Estes et al. 2010) | 2010 | ? | ? | ? | NG | 37C (24hr), 25C (24hr) | ? |
| (Dunbar et al. 2012) | 2012 | ? | ? (O.N.) | ? | NG | 37C (24hr), 25C (24hr) | ? |
| (Pukkila-Worley et al. 2012) | 2012 | LB | 37C (14-15hr, 3mL) | 10 µL | NG +FUDR +1%DMSO +RPW-24 | 37C (24hr), 25C (24hr) | 25C |
| (Feinbaum et al. 2012) | 2012 | LB | 37C (16hr, 150 µL) | 10 µL (6-well plate) | NG | ? | 25C |
| (Kirienko et al. 2014) | 2014 | LB | 37C (12-16h, 5mL) | 10 µL | NG +FUDR | 37C (24hr), 25C (24hr) | 25C |
| (Pellegrino et al. 2014) | 2014 | ? | ? (O.N.) | ? | NG | 25C (O.N.), 37C (24hr) | 25C |
| (Zou et al. 2014) | 2014 | ? | ? | 100 µL | NGM +FUDR | 37C (24hr), 25C (24hr) | 25C |
| (Meisel et al. 2014) | 2014 | LB | 37C (O.N., 2-3mL) | 7 µL or 15 µL | NG +FUDR | 37C (O.N.), Room temp. (1 day) | 25C |
| (Hilbert and Kim 2017) | 2017 | LB | 37C (O.N., 3mL) | 7 µL | NG | 37C (O.N.), Room temp. (2 days) | ? |
| (Lee and Mylonakis 2017) | 2017 | LB | ? (Until OD600 2.5, 4mL) | ? | NG +FUDR | 37C (24hr), 25C (8-16hrs) | ? |
| (Dinić et al. 2021) | 2021 | LB | 37C | ? | NG +FUDR | 37C (24hr), 25C (48hr) | 20C |
| (de Lucas et al. 2021) | 2021 | King's broth | ? (O.N.) | 10 µL | NG | 37C (O.N.) | ? |
| (Ambreetha and Balachandar 2022) | 2022 | ? | ? (O.N.) | 50 µL OD 0.5 | NGM | ? | 20C |
| (Hoang et al. 2024) | 2024 | LB | 30C (O.N.) | ? | NGM | 30C (24hr) | 20C |
| (Xiao et al. 2024) | 2024 | LB | 37C (O.N.) | ? | NGM | - | 25C |
| (Liu et al. 2024) | 2024 | LB | 37C (16hrs, 5mL) | 150 µL | NGM +FUDR | 37C (24hr), 25C (9-13hr) | 25C |
| (Gainey et al. 2024) | 2024 | LB? | 37C (14 or 18 hrs) | 750 µL OD 1.0 | NGM | 25C (48hrs) | 20C |
| (Xue et al. 2024) | 2024 | LB | 37C (16hrs) | 250 µL | NGM | ? | 25C |
| (Marogi et al. 2024) | 2024 | ? | O.N. | OD 1.0 | NGM | 37C (O.N.), 25C (Briefly) | ? |
| [this paper] | 2026 | LB | 30C (O.N., 5mL) | 100 µL OD 0.05 | NG | 30C (24hrs), 25C (24hrs) | 25C |

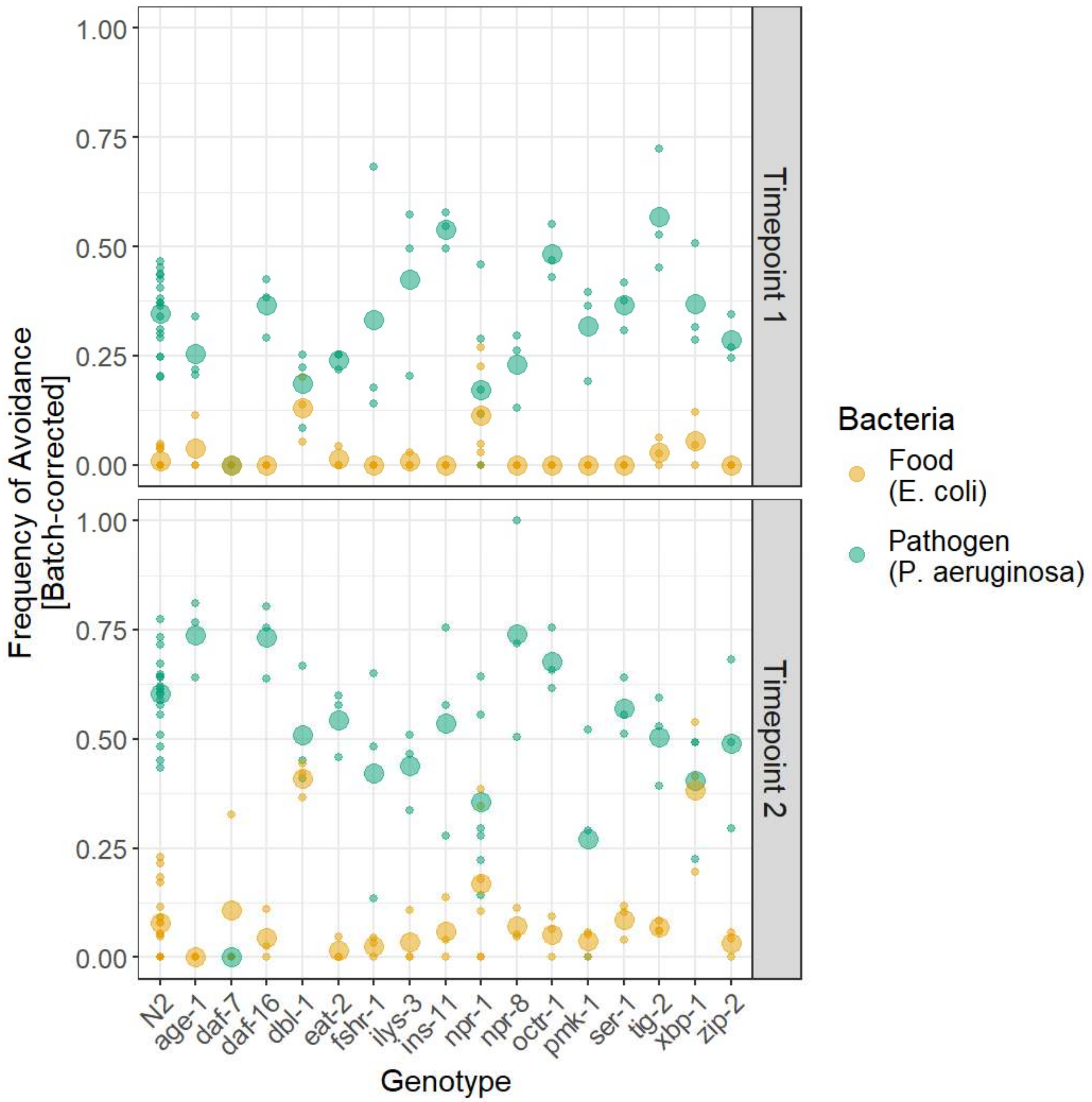


**Figure S1. Host genotypes vary in behavioral avoidance of bacteria.** Wild-type (N2) and mutant genotypes with single gene knockouts exposed to a spot of food or pathogen bacteria on an otherwise-empty agar plate for 24 hours (Timepoint 1) and 48 hours (Timepoint 2), then the number of hosts on the spot versus avoiding the spot was counted. Small points denote individual replicates while large points denote the average.

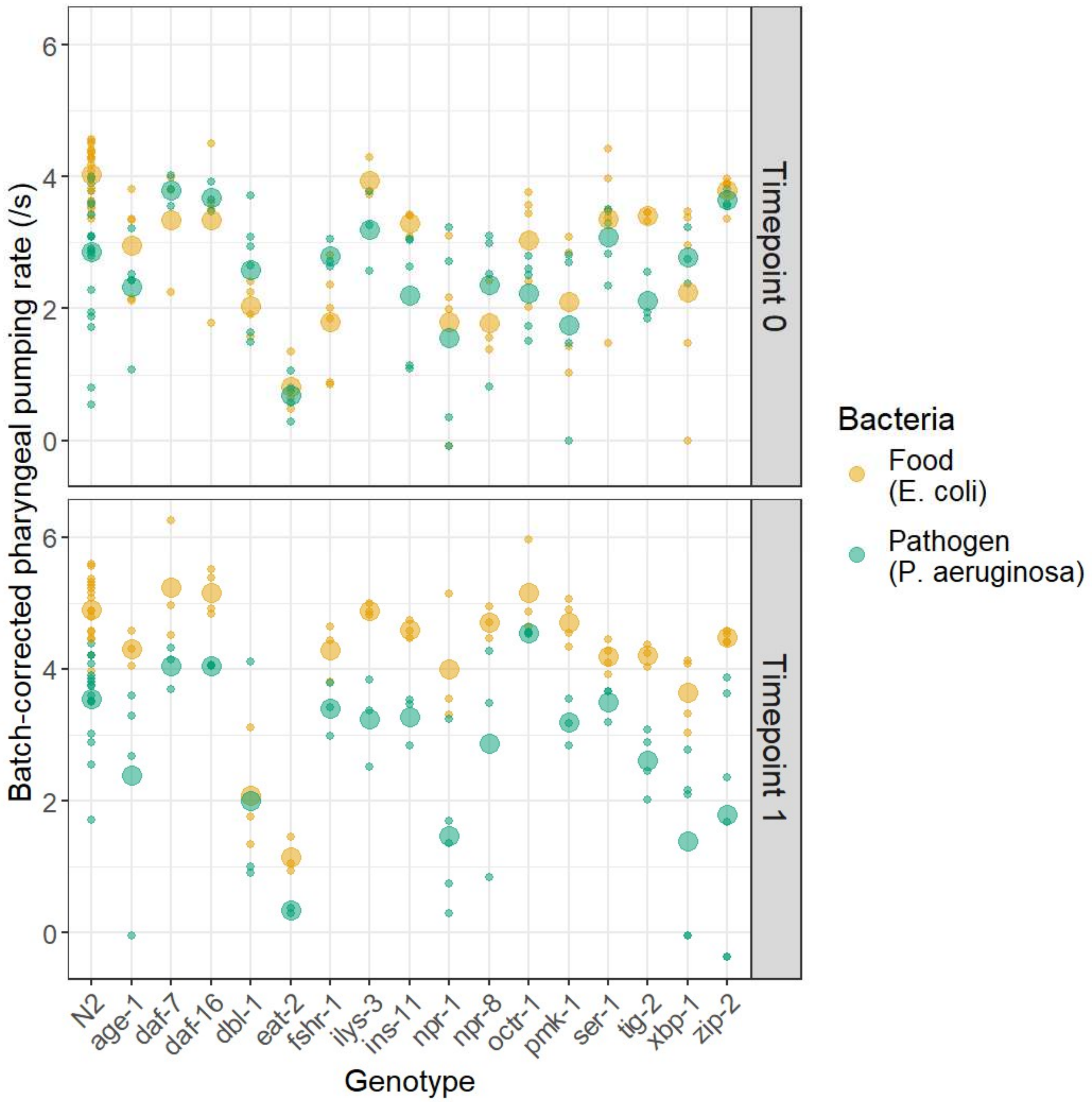


**Figure S2. Host genotypes vary in uptake of bacteria.** Wild-type (N2) and mutant genotypes with single gene knockouts were exposed to lawns of food or pathogen bacteria for 0 hours (Timepoint 0) or 24 hours (Timepoint 1), at which point the pharyngeal muscle movements of at least 3 worms per strain were manually counted from 30s videos. Small points denote individual replicates while large points denote the average.

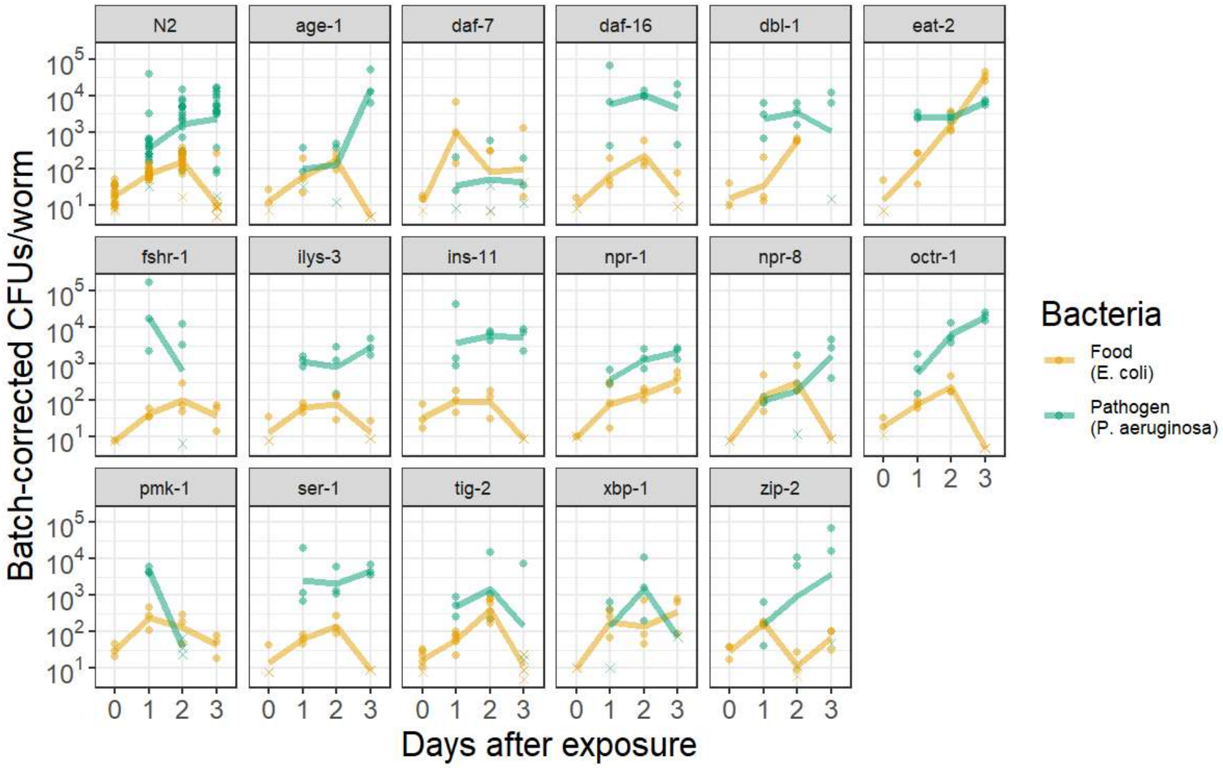


**Figure S3. Host genotypes vary in susceptibility to colonization by bacteria.** Wild-type (N2) and mutant genotypes with single gene knockouts were exposed to lawns of food or pathogen bacteria for three days. 10 worms of each genotype were picked from each of three replicate plates for each strain in each condition and at each timepoint. After picking, the exterior of the worms was sterilized before the gut contents were plated to determine the average colony-forming units (CFUs) per worm. Small points denote individual replicates while lines denote the average, 'x's denote replicates where the colony forming units were below the limit of detection, and are plotted at the limit of detection for that replicate.

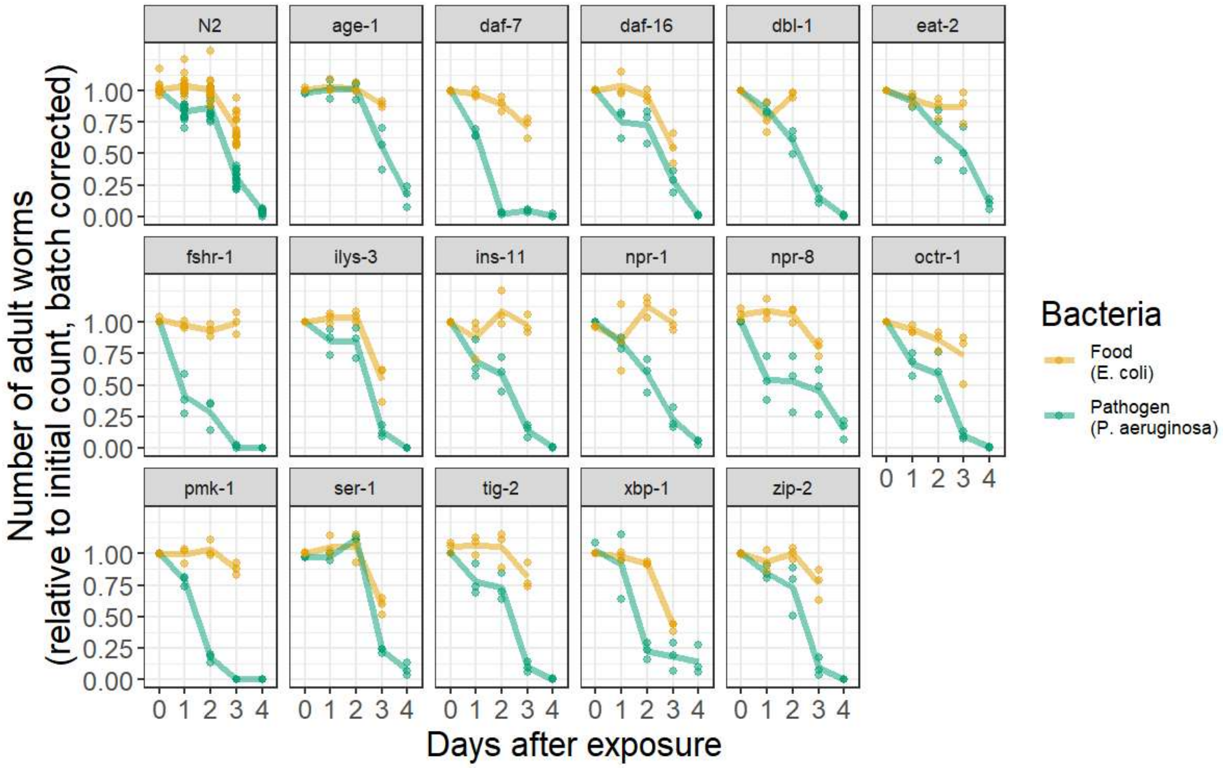


**Figure S4. Host genotypes vary in mortality during exposure to bacteria.** Wild-type (N2) and mutant genotypes with single gene knockouts were exposed to lawns of food or pathogen bacteria for three or four days, respectively. Three replicate plates were counted for each genotype in each condition. Small points denote individual replicates while lines denote the average. dbl-1 on food plates was not counted at 3 days because the next generation of nematodes had reached adulthood.

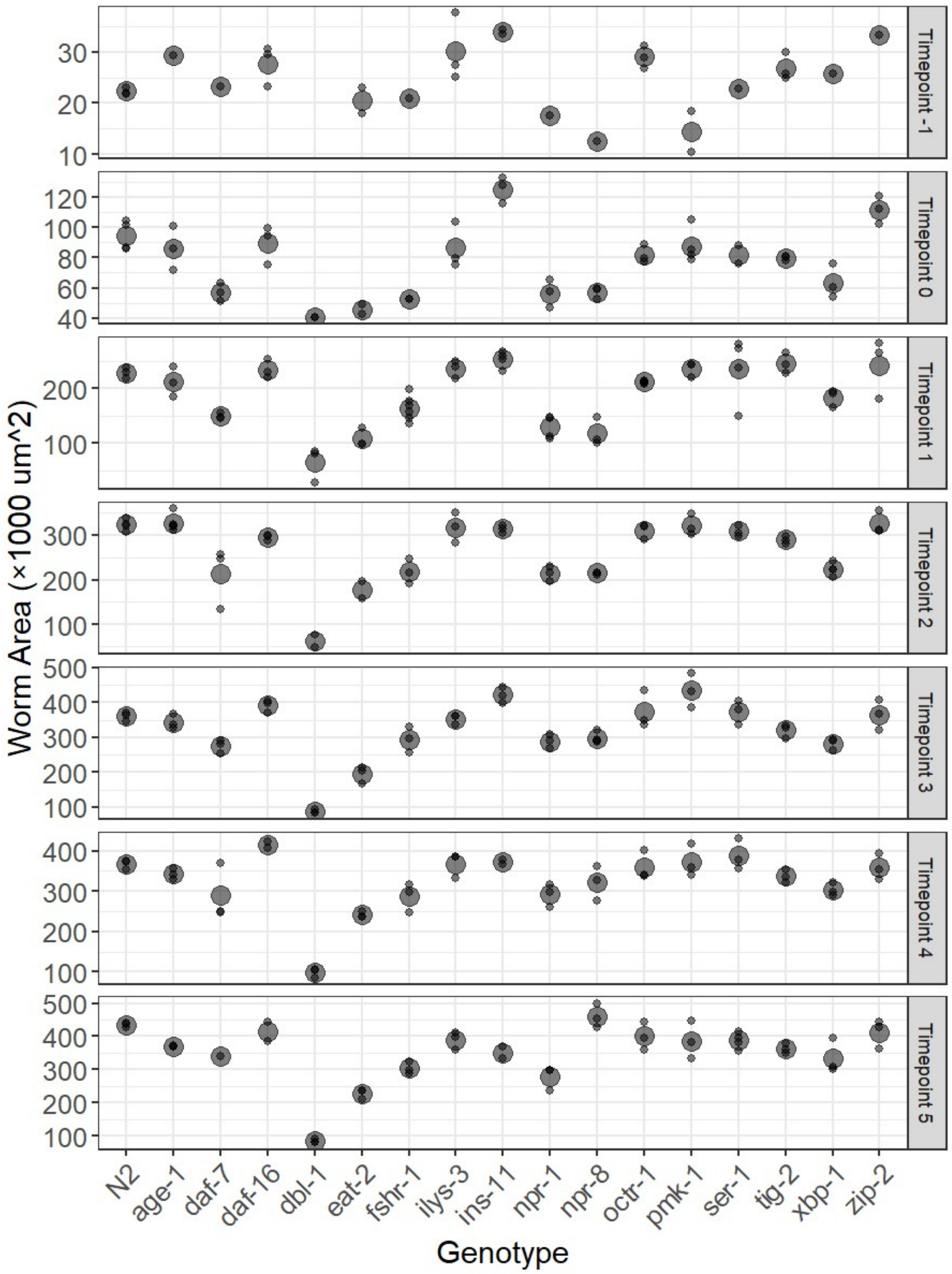


**Figure S5. Host genotypes vary in projected body area.** Wild-type (N2) and mutant genotypes with single gene knockouts were reared on food bacteria for seven days (with Timepoint 0 corresponding to 48 hours after plating, when worms are typically transferred to exposure conditions). Each day, worms were imaged (n ≥ 3 each genotype). After imaging, the projected body area of each worm was calculated using WormLab. Small points denote individual replicates while large points denote the average. Videos of dbl-1 at Timepoint -1 (i.e. 24 hours after plating) did not have sufficient resolution for WormLab to accurately quantify worm size, and so were excluded.

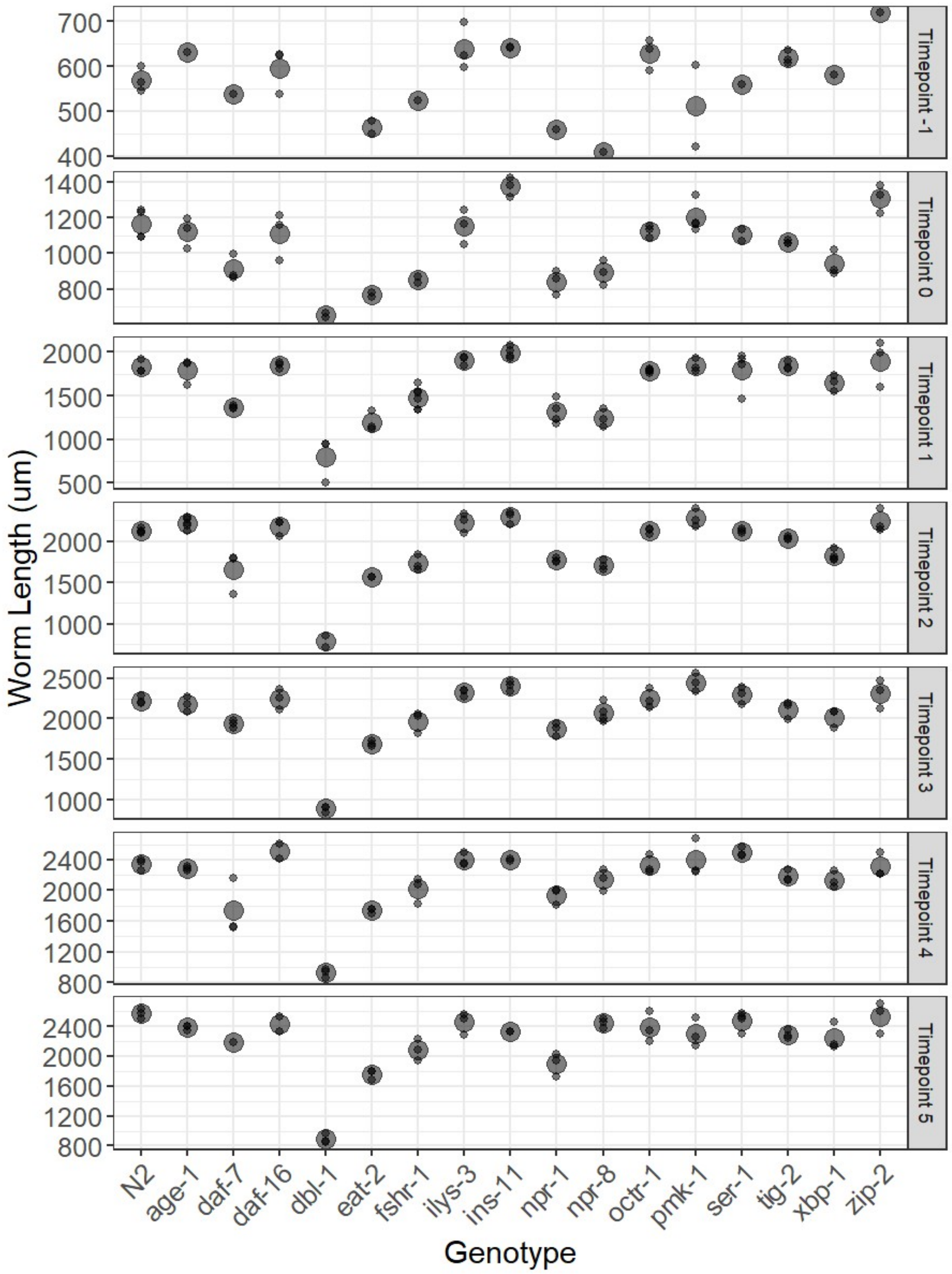


**Figure S6. Host genotypes vary in body length.** Wild-type (N2) and mutant genotypes with single gene knockouts were reared on food bacteria for seven days (with Timepoint 0 corresponding to 48 hours after plating, when worms are typically transferred to exposure conditions). Each day, worms were imaged (n ≥ 3 each genotype). After imaging, the length of each worm was calculated using WormLab. Small points denote individual replicates while large points denote the average. Videos of dbl-1 at Timepoint -1 (i.e. 24 hours after plating) did not have sufficient resolution for WormLab to accurately quantify worm size, and so were excluded.

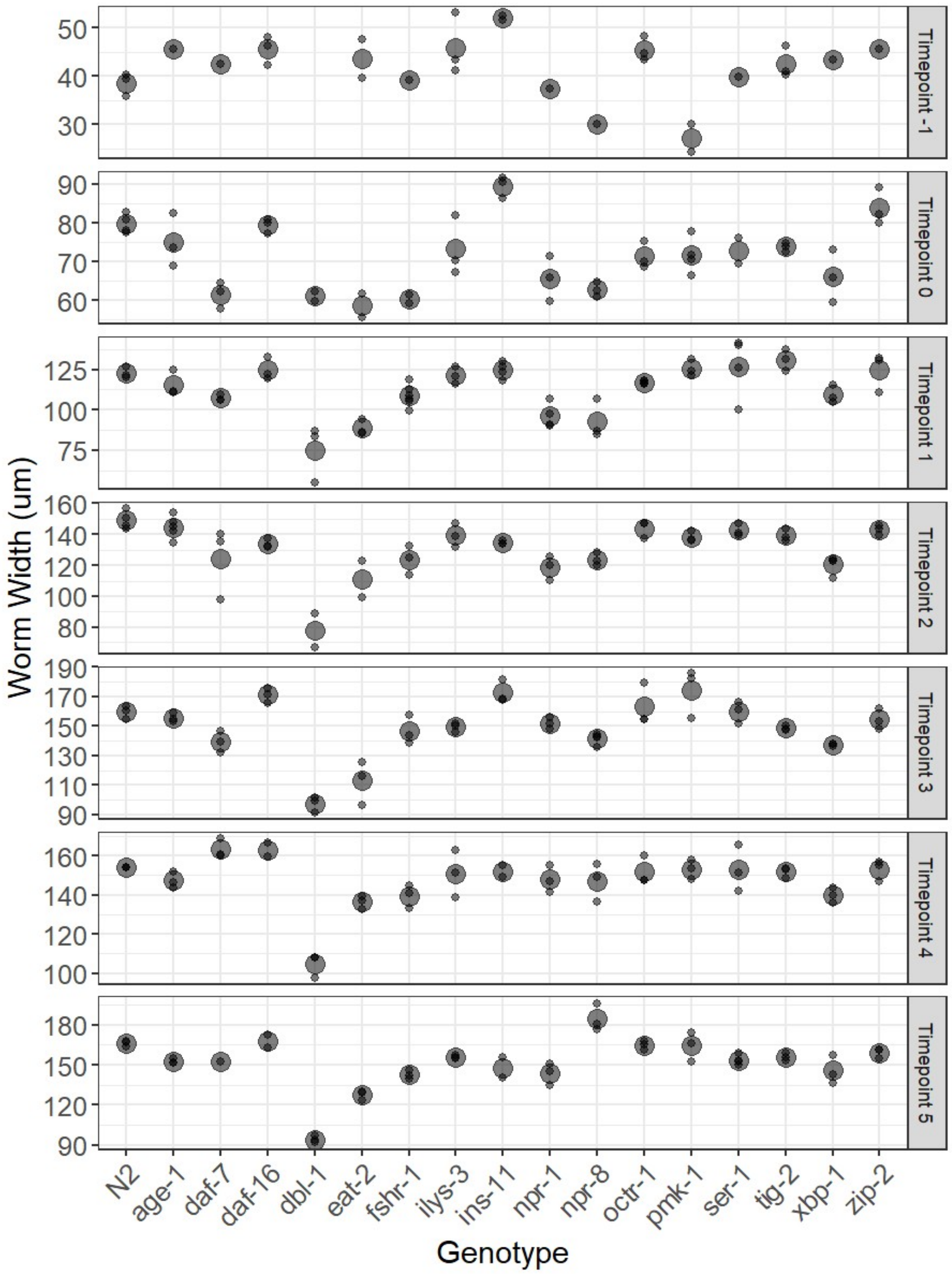


**Figure S7. Host genotypes vary in body width.** Wild-type (N2) and mutant genotypes with single gene knockouts were reared on food bacteria for seven days (with Timepoint 0 corresponding to 48 hours after plating, when worms are typically transferred to exposure conditions). Each day, worms were imaged (n ≥ 3 each genotype). After imaging, the width of each worm was calculated using WormLab. Small points denote individual replicates while large points denote the average. Videos of dbl-1 at Timepoint -1 (i.e. 24 hours after plating) did not have sufficient resolution for WormLab to accurately quantify worm size, and so were excluded.

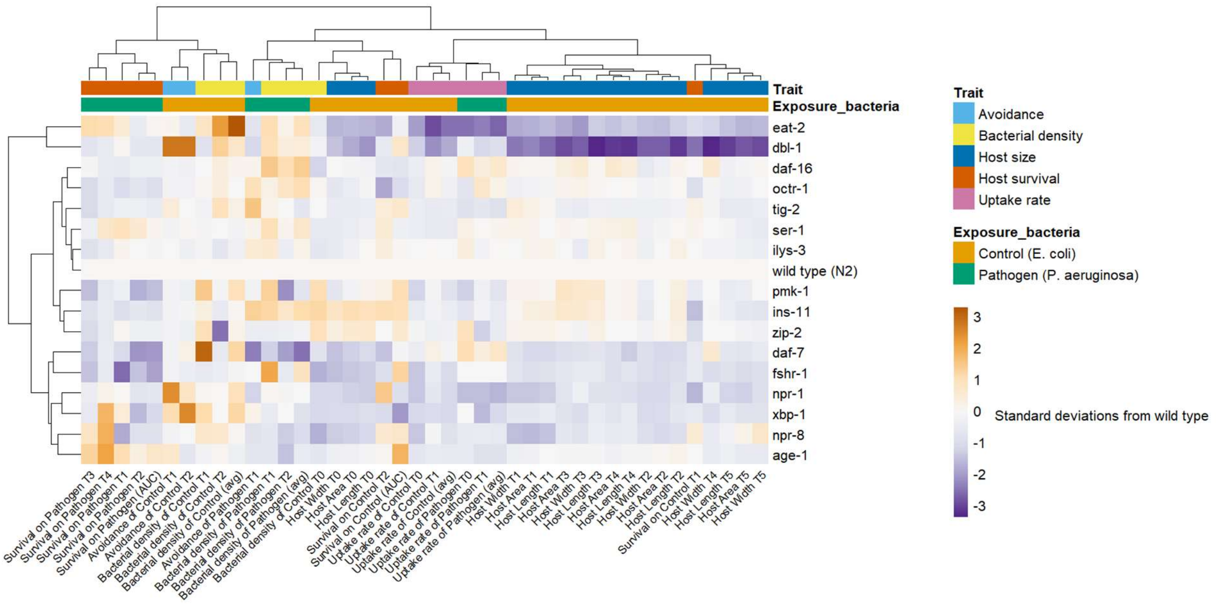


**Figure S8. Host traits cluster across timepoints.** Average values for each genotype from Figs S1-S7 were replotted after being centered at the wild type value and scaled proportional to the standard deviation among different genotypes. Clustering of traits and genotypes was done by mean-centering and scaling the traits, then clustering either traits or genotypes with Euclidean distance and Ward's method for agglomeration.

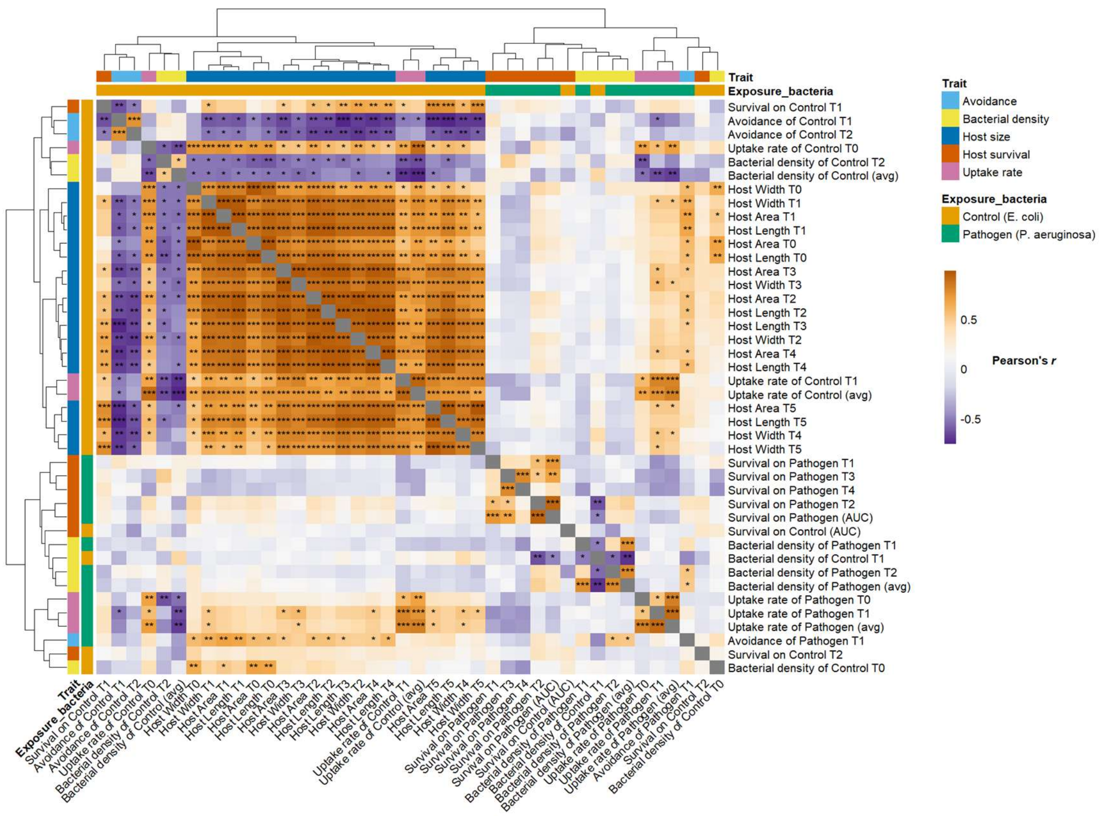


**Figure S9. Host traits cluster across timepoints, and some form a highly correlated cross-trait cluster.** Data from Fig S8 was used to calculate the correlation of each trait against each other trait among the 17 host genotypes. Clustering of traits was done by Euclidean distance and complete linkage of the absolute value of the correlation coefficient. Asterisks denote the two-tailed significance of Pearson's *r*: *** for p < 0.001, ** for p < 0.01, and * for p < 0.05.
